# Using very high-resolution satellite imagery and deep learning to detect and count African elephants in heterogeneous landscapes

**DOI:** 10.1101/2020.09.09.289231

**Authors:** Isla Duporge, Olga Isupova, Steven Reece, David W. Macdonald, Tiejun Wang

**Author notes:** Equal contribution (Isla Duporge, Olga Isupova).

## Abstract

1. Satellites allow large-scale surveys to be conducted in short time periods with repeat surveys possible <24hrs. Very high-resolution satellite imagery has been successfully used to detect and count a number of wildlife species in open, homogeneous landscapes and seascapes where target animals have a strong contrast with their environment. However, no research to date has detected animals in complex heterogeneous environments or detected elephants from space using very high-resolution satellite imagery and deep learning.
2. In this study we apply a Convolution Neural Network (CNN) model to automatically detect and count African elephants in a woodland savanna ecosystem in South Africa. We use WorldView-3 and 4 satellite data – the highest resolution satellite imagery commercially available. We train and test the model on eleven images from 2014-2019. We compare the performance accuracy of the CNN against human accuracy. Additionally, we apply the model on a coarser resolution satellite image (GeoEye-1) captured in Kenya to test if the algorithm can generalise to an elephant population outside of the training area.
3. Our results show the CNN performs with high accuracy, comparable to human detection capabilities. The detection accuracy (i.e., F2 score) of the CNN models was 0.78 in heterogeneous areas and 0.73 in homogenous areas. This compares with the detection accuracy of the human labels with an averaged F2 score 0.77 in heterogeneous areas and 0.80 in homogenous areas. The CNN model can generalise to detect elephants in a different geographical location and from a lower resolution satellite.
4. Our study demonstrates the feasibility of applying state-of-the-art satellite remote sensing and deep learning technologies for detecting and counting African elephants in heterogeneous landscapes. The study showcases the feasibility of using high resolution satellite imagery as a promising new wildlife surveying technique. Through creation of a customised training dataset and application of a Convolutional Neural Network, we have automated the detection of elephants in satellite imagery with as high accuracy as human detection capabilities. The success of the model to detect elephants outside of the training data site demonstrates the generalisability of the technique.

## 1. Introduction

The planet is in the geological era of the Anthropocene, during which human activity is the driving force of change. Many wildlife species are under threat across their geographical range as we are currently undergoing the sixth-mass extinction [1-3]. Reliable, accurate and up-to-date data on wildlife numbers is essential to monitor population fluctuations and identify causes of decline. Various methods are used for conducting population counts e.g. line transect surveys [4], dung and track counts [5], bio-acoustic monitoring [6], camera trap [7] and aerial surveys [8], among others.

Satellite remote sensing has recently emerged as a new viable monitoring technique for detecting wildlife. It has been used to successfully identify and count a number of wildlife species in open, homogeneous landscapes and seascapes. The benefits of this monitoring technique are numerous; large spatial extents can be covered in short time periods making repeat surveys and reassessments possible at short intervals. For example, the satellite used in this study, Worldview-3 has an average revisit time of less than one day. It is capable of collecting up to 680,000 square kilometres every 24hrs. Satellite images are captured over large areas in one shot so issues with double counting and miscounts are largely eliminated. Satellite remote sensing is unobtrusive as it requires no human presence eliminating the risk of disturbing the species being surveyed. This remains a key concern in other surveying techniques [9]. Image acquisition is automated and less logistically complicated compared with traditional aerial surveys [10] and setting up camera trap grids or audio loggers. Censuses can be carried out without concern for human safety providing an ability to survey previously inaccessible areas. For example, in the case of the Emperor penguin, new colony locations were detected in a pan-continental survey of the Antarctic coast [7, 13]. Additionally, cross border areas can be surveyed without requiring multiple national civil aviation permissions.

Detecting wild animals in satellite imagery is influenced by body size, background complexity and contrast between species and surrounding habitat. Seascapes provide a uniform high contrast background context against which whales have been identified in known breeding, calving and feeding grounds [11-13] and flamingos have been identified on a lake [14]. Spectrally uniform rocky outcrops and ice have been used to identify several marine species i.e. Emperor and Adelie penguins [15-19], Elephant and Weddell seals [20, 21], Masked Boobies [22] and Albatross [23]. Several Arctic species have been identified against snow using shadow and body contrast for detection i.e. Polar bears [10, 24-26] and muskoxen [25]. On the African continent only two studies have detected mammals (in the case of wildebeest and zebra) using satellite imagery in open savannah [27, 28]. Detection has only been tested in these homogeneous monochrome environmental contexts. No study has yet, to the best of our knowledge, detected species in complex heterogeneous landscapes from space.

Various methods have been used to detect species in satellite imagery. The majority of studies have manually counted species in imagery using several observers for cross-validation. However, manual methods are unfeasible if large areas are surveyed, as counting is labour and time intensive and counts tend to be error-prone [29]. Several studies have relied on environmental proxies and indirect ecological signs of animal presence e.g. burrows [30], mounds [31], changes in vegetation from nest signatures [22] and faecal stains in the case of penguins [15-19, 32]. Image differencing is a technique where satellite images are captured in the same location at different times. This technique is used for environmental change detection [33] e.g. deforestation and land use change [34-36], identification of fire [35, 37], droughts [38, 39] or floods [40, 41]. Three studies used short-time image differencing to detect polar bears from space [10, 24, 26]. Image differencing is possible in cases where multi-temporal imagery is available, and species can be differentiated from static objects. e.g. rocks. Images can be acquired via targeted satellite tasking on specific days; however, this is more costly than using archive imagery. Cloud cover, environmental variability and changing sea states can impede ground visibility which is problematic when image differencing and tasking is used.

Several studies have applied a form of supervised or semi-supervised classification approaches to detect species in satellite imagery. One form of image segmentation using semi-supervised classification is thresholding method. Pixel values are classified relative to a set of threshold values that distinguish species from background. Thresholding method does not make use of geometric information but rather relies on spectral signatures (pixel value combinations). Thresholding method is reliant on the human classifier to set accurate thresholds which is helped by familiarity with the species and environment [27]. This technique is effective in homogeneous environments where species have strong spectral separability from background context. However, in cases where pixel values of species and background context are similar it is difficult to draw accurate distinctions.

The introduction of Convolutional Neural Networks (CNN) in machine learning has revolutionised the field of computer vision since 2012 [42]. Machine learning has become a new essential tool used by ecologists to detect wildlife in imagery e.g. camera trap images, aerial survey images, and unmanned aerial vehicle (UAV) images [43-48]. However, automated detection of wildlife from satellite imagery is still in its infancy. To the best of our knowledge only two species have been detected in satellite imagery using deep learning: albatross [49] and whales [50, 51]. Object detection applications are now easier to develop than ever before. High-performance off-the-shelf solutions have made machine learning solutions accessible to non-specialists. These techniques can now leverage massive image datasets e.g. ImageNet (>14 million images across 20,000 classes) obtaining significant performance improvements compared to previous methods based on manually engineered features [52]. A Convolutional Neural Network (CNN) is a deep learning artificial neural network architecture that has been extensively used for object detection and recognition in recent years. The ‘deep’ stems from the use of multiple layers in the network. In this study we test whether it is possible to detect the world’s largest terrestrial mammal – the African elephant – using deep learning via a CNN.

The population of African elephants (*Loxodonta africana*) has plummeted over the last century due to poaching, retaliatory killing from crop raiding and habitat fragmentation [53-55]. To ensure conservation is achieved accurate monitoring is vital. Inaccurate counts can result in misallocation of scarce conservation resources and misunderstanding population trends. Existing techniques are prone to miscounting. The most common survey technique for elephant populations in savannah environments is aerial counts from manned aircraft [8]. Aerial surveys are conducted either as total counts – flying closely spaced transects, or sample counts, covering 5-20% of an area and extrapolating to a larger area. Several studies have shown that observers on aerial surveys often miscount due to fatigue and visibility issues resulting in over-estimates [56-58]. Aerial surveys can be costly, logistically challenging in terms of finding suitable runways and refuelling stops and time consuming in terms of getting appropriate permissions. This is particularly relevant in cross-border areas where multiple national permissions are required. Remotely sensing elephants using satellite imagery and automating detection via deep learning may provide a novel avenue for surveying while mitigating several of the challenges outlined above.

In this study we investigate the feasibility of using very high-resolution satellite imagery to detect wildlife species in heterogenous environments with deep learning. To test this, we use a population in Addo Elephant National Park, South Africa where herds move between open savannah habitat and closed heterogeneous woodland and thicket.

## 2. Methods

### 2.1. Study Site

Addo Elephant National Park in South Africa was chosen as the study site. It provides a spectrally complex heterogeneous background with a high concentration of elephants. The park is the third largest park in South Africa at 1,640 km^2^. Different areas of the park have been sectioned off for conservation purposes - elephants were identified in the Main Camp section of the park surrounding Hapoor Dam (Figure 1). The Main Camp is a combination of dense shrubland and low forest (e.g. porkbush (*Portulcaria afra)*, White milkwood *(Sideroxylon inerme)*, Cape leadwort (*Plumbago auriculate*) and open grassland [59, 60]. Over six hundred elephants move between these habitats [61, 62]. Elephants cover themselves in mud to cool down and adopt a range of postures when foraging, playing, sleeping [63, 64] so their shape and colour is continually changing. The background environment is also changing as they move between open savannah and woodland and take shelter under trees in the mid-day sun. The satellite is on a sun synchronous orbital path, so satellite images of the study area are captured between 10.10-10.45am local time. The bright morning light improves image clarity, as elephants gather at water holes in the morning which makes them easy to identify (Figure 1). The park has rainfall year-round [65] and four seasons can be broadly delineated as early wet season (Oct -Dec), late wet season (Jan-March), early dry (Apr-Jun) and late dry season (July-Sept) [61, 62].

**Figure 1.**
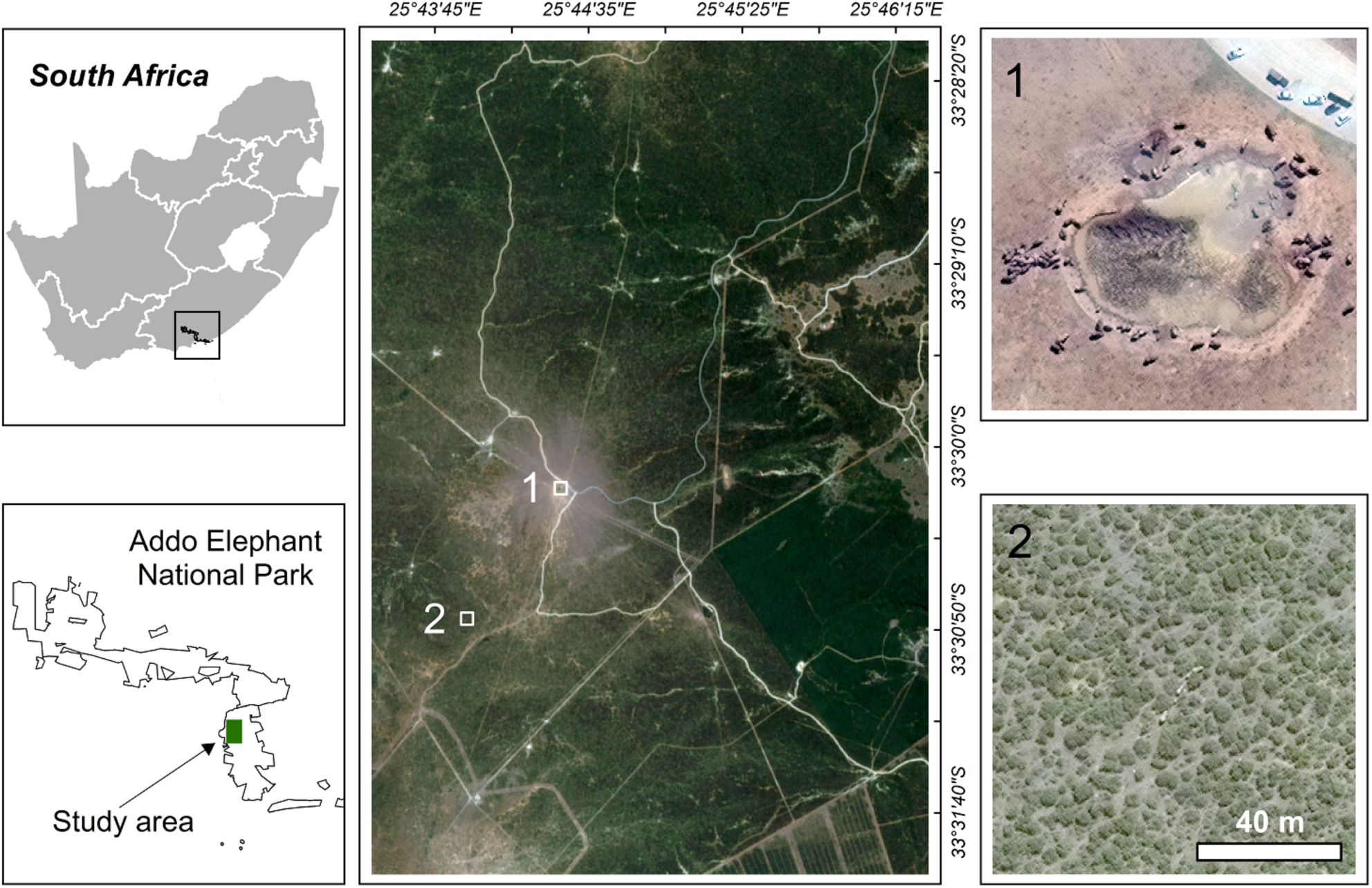
Location of the study area in - Addo Elephant National Park, South Africa. Two example WorldView-3 images showing 1) Elephants in open homogeneous area around Hapoor Dam, 2) Elephants in heterogenous woodland and thicket area. Satellite image (c) 2020 Maxar Technologies

To ensure a representative and diverse sample of elephants in the park we include training and test labels from images captured in different seasons and years (Table 1).

**Table 1.** List of satellite images used in the training and test dataset

| *Date of acquisition* | *Satellite* | *Elephant labels in training dataset* | *Elephant labels in validation dataset* | *Elephant labels in test dataset* |
| --- | --- | --- | --- | --- |
| 01_12_2014 | WV3 | 197 | 52 | 10 |
| 29_01_2016 | WV3 | 178 | 9 | 11 |
| 10_02_2016 | WV3 | 259 | 19 | 32 |
| 03_04_2017 | WV3 | 26 | 19 | 5 |
| 22_11_2017 | WV4 | 10 | / | / |
| 11_01_2018 | WV4 | 117 | / | 23 |
| 27_03_2018 | WV4 | 236 | 16 | 24 |
| 08_10_2018 | WV4 | 119 | 1 | 59 |
| 20_01_2019 | WV3 | 22 | / | / |
| 11_08_2009 | GeoEye-1 | / | / | 32 |

### 2.2. Dataset generation and satellite image pre-processing

WorldView-3 and WorldView-4 are the highest resolution satellite imagery commercially available. They provide imagery at 31cm resolution - WorldView-4 is now out of action but two years of archive imagery is available. The image archive for all WorldView 3 & 4 satellite images from Maxar Technologies (formerly DigitalGlobe) was searched via the Secure Watch Platform [https://www.digitalglobe.com/products/securewatch]. We restricted the search to images that contain less than 20% cloud cover and acquired less than 25% off-nadir (degree off centre of image captured). We selected eleven images that met these specifications between 2014-2019.

Each image was downloaded in two formats: orthorectified images in natural colour and orthorectified panchromatic image. We processed the images using a pan-sharpening algorithm from ERDAS IMAGINE software package (ERDAS, Inc., Atlanta, GA, USA) Pan-sharpening is an automatic image fusion process that uses the multispectral bands red (620–750 nm), green (495– 570 nm), blue (450–495 nm) at 1.24 m resolution and the higher resolution panchromatic band at 31 cm to produce a high-resolution multispectral image. We tested several pan-sharpening algorithms - the Gram-Schmidt pan-sharpening algorithm provided the cleanest visual result and was applied to all images. The satellite images were converted so that pixel values were in the range of [0, 255] and the images were sliced into 600×600pixel sub images to make them compatible with the deep learning software.

### 2.3. Labelling training data in satellite images

The images were visually scanned for elephants before sub-setting into smaller areas where we identified congregations of elephants. In total, 1125 elephants were identified in the training image dataset. To ensure training labels are representative of elephants at different times, images were selected for different seasons and years in both closed i.e. dense shrubland and forest. and open areas of the park i.e. grassland and bare land. Images were labelled by defining bounding boxes around each individual elephant using the graphical image annotation tool LabelImg [https://github.com/tzutalin/labelImg] [66] shown in Figure 2.

**Figure 2.**
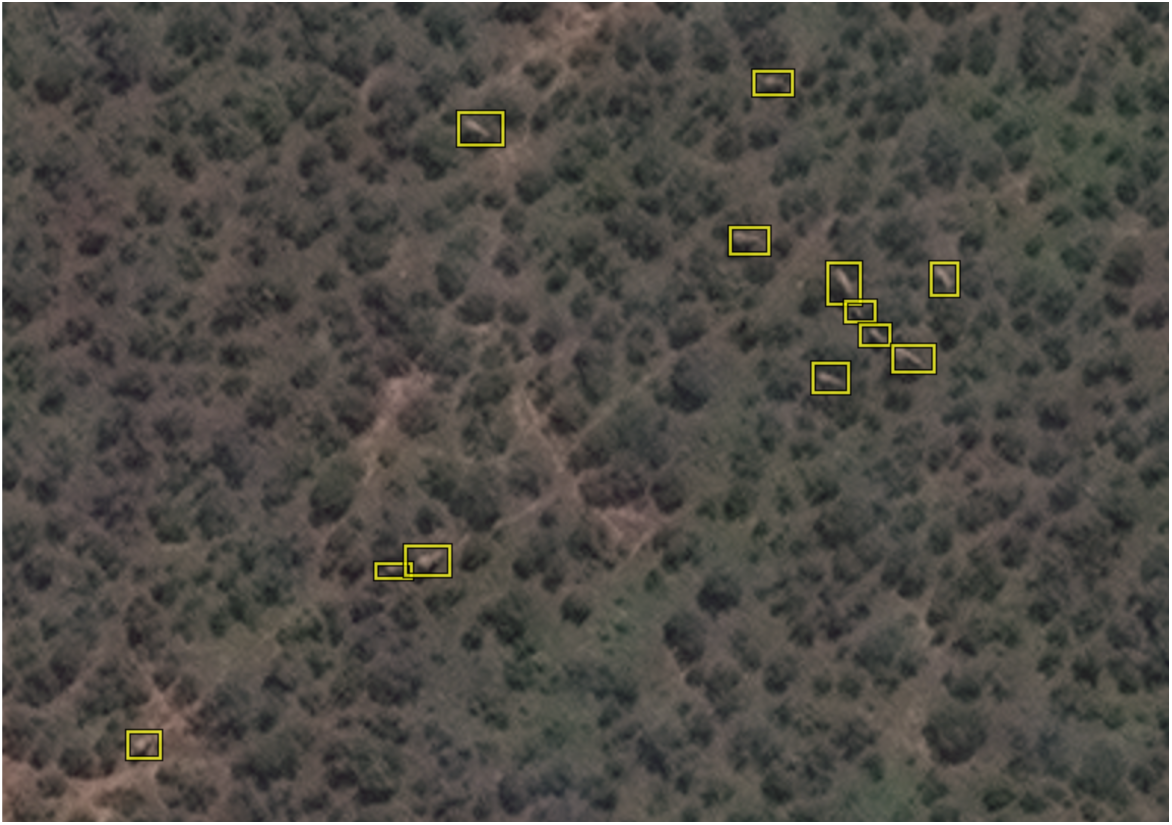
Example of elephant labels in a heterogenous area, Addo Elephant National Park, South Africa. Satellite image (c) 2020 Maxar Technologies

The baseline we deem as the true number of elephants is a labelled dataset doubled screened by two annotators – an Ecologist and Machine Learning Scientist. Any ambiguous labels that were not identified by both annotators were removed. We use the method of comparing the accuracy of detections from human volunteer annotators and CNN performance against this baseline control count [46, 67]. The same test images used to evaluate CNN performance were sent to 51 human volunteer annotators. The images were labelled by the volunteers using the VGG Annotation Tool [http://www.robots.ox.ac.uk/~vgg/software/via/][68]. Volunteer annotators represent a cross-section of machine learning scientists, biologists, general public and park rangers who work with elephants in Southern Africa. The labellers vary in terms of computer literacy and familiarity with elephant behaviour and habitat preference. The experiment involving human participants was approved by the University of Oxford CUREC Ethical Committee [R64699].

### 2.4. Training and validating the Convolutional Neural Network model

A CNN is a feed-forward neural network designed to process large-scale images by considering their local and global characteristics [69]. A neural network is typically comprised of multiple layers connected by a set of learnable weights and biases [70]. Convolutional layers represent a set of filters, each able to identify a particular feature in the image. These filters are fed small image patches whilst they scan across the image and generate feature maps for analysis by the next layer. The CNN comprises an alternating sequence of convolutional and pooling layers. Pooling layers are used to reduce the dimensionality of the feature maps to improve computational efficiency. Nonlinear activations are stacked between convolutional layers to enrich their representational power [71]. The last layer of the network is fully connected and performs classification [72]. Convolutional neural networks have become a key tool in image classification. They are now comparable to human performance in a number of challenging image classification tasks e.g. face verification, various medical imaging tasks [73, 74].

We used the TensorFlow Object Detection API [https://github.com/tensorflow/models/tree/master/research/object_detection] to build our model [75]. This API provides implementations of different deep learning object detection algorithms. In a preliminary assessment of the models available, we selected the model referred to as faster_rcnn_inception_resnet_v2_atrous_coco as it provided the best result and it was used for all the experiments presented. This model is a Faster Region CNN (RCNN) model – after layers that are used to extract features there is a subbranch to propose regions that may contain objects and a subbranch that predicts the final object class and bounding box for each of these regions [76]. The model we used has an Inception ResNet [77] backbone – this is the underlying CNN that is used for feature extraction. We used the model pretrained on the Common Objects in Context (COCO) dataset for object detection [https://cocodataset.org/] [78]. We used default values for hyperparameters of this model from the API.

Training a CNN requires images to be split into training, validation and test sets. In total, 188 sub images from nine different satellite images were used for training. These training images contain 1270 elephant labels of which 1125 are unique elephants. There is an overlap of 50 pixels between sub images, when elephants appear at the edge of one sub image the overlap ensures they appear in whole on the neighbouring sub image. Twelve sub images containing 116 elephant labels were left out as a validation dataset. The validation dataset is used to tune the hyperparameters, to define the confidence threshold (above which predictions from the model are counted as detections) and to identify the optimal length of CNN training (see Figure 3).

**Figure 3.**
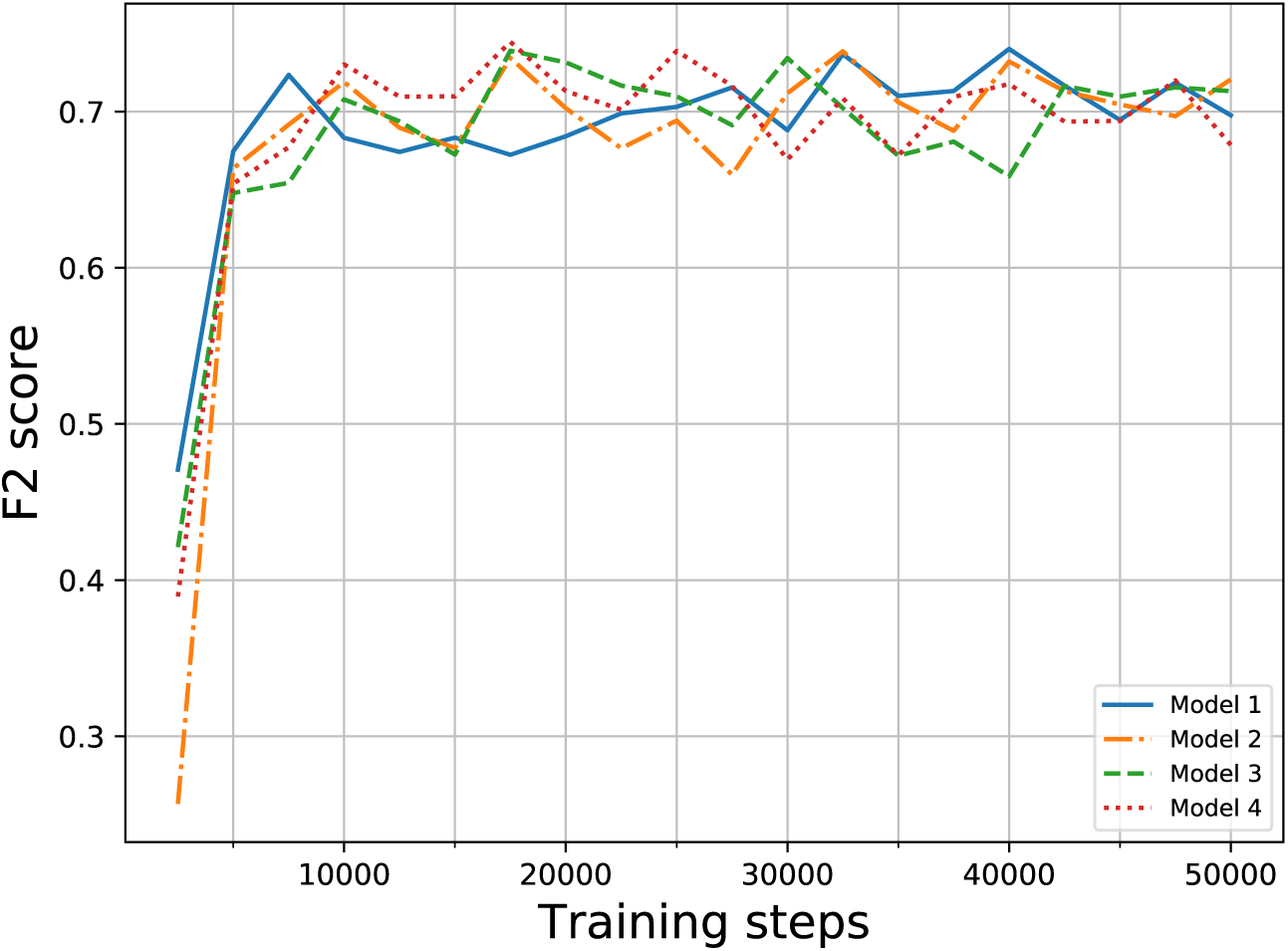
F2 score obtained by each of the four models considered over training steps on the validation dataset. All models converged during the first 50,000 training steps

### 2.5. Test dataset

The test dataset used to test the CNN against human annotator performance contains 164 elephants across seven different satellite images. These images do not overlap with any of the training or validation sub images. The test sub images cover both heterogeneous and homogeneous areas of the park from different seasons and years (see Table 1).

In addition, an image from the Maasai Mara in Kenya was used to test the transferability of the CNN under broader image conditions. The image comes from Geoeye-1 a lower resolution satellite (41cm) compared to the images used for training which come from Worldview 3 & 4 (31cm). This image allowed us to test the generalisability of our algorithm to a different environment and satellite.

### 2.6. Accuracy assessment

We compare the accuracy of detections from the human volunteer annotators and CNN against our count which we deem as the baseline i.e. the true number of elephants in the images. To calculate the deviation from this baseline we generate an F2 score.

Object detection performance is usually evaluated by *precision* and *recall*:

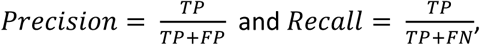

where *TP* stands for true positives (correctly predicted elephants), *FP* stands for false positives (predicted elephants that are not actually elephants, also called false detections), *FN* – false negatives (elephants that are not detected by the model, also called missed detections).

The CNN gives an output in the form of bounding boxes, the same format we use for the training labels. We count any intersection between predicted and true bounding boxes as a true positive (i.e. the intersection over union threshold used to determine correct predictions was set to 0). Human volunteer annotators provide point detections for elephants. If these point detections were inside true bounding boxes, they were counted as true positives.

In precision and recall both types of errors – false positives and false negatives – are weighted equally. However, as it is more time consuming for a human annotator to check an entire image for missing elephants (false negatives) as compared with reviewing detected elephants and eliminating false positives we decided to use an F2 (*F*_*β*_ with *β* = 2) score. The F2 combines precision and recall in such a way that more emphasis is put on false negatives [79, 80]:

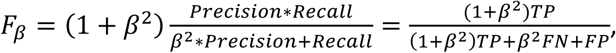

which for *β* = 2 is equivalent to

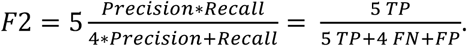

Performance of object detection algorithms are often measured by *average precision* [75] i.e. the area under a precision-recall curve that is obtained by varying the threshold of the confidence score. This threshold determines which of the predicted bounding boxes are considered as final detections. Average precision allows comparison between different algorithms without the need to specify this threshold. Since our goal was to compare the algorithm performance with human performance and humans did not provide a confidence score for their detections, we could not use this metric.

The training process is stochastic due to the stochastic gradient descent algorithm used for optimisation of neural network weights. We ran the CNN four times to explore how stable the algorithm output is with respect to the stochastic training process. Neural networks models are commonly run as many times as time and availability of computational resources allow. Each of the models ran for 50,000 training steps (i.e. the number of times the weights were updated by the gradient descent algorithm) on the training dataset and the performance was evaluated on the validation dataset every 2,500 training steps (Figure 3). All the models reached a plateau in F2 score after around 10,000 training steps on the validation dataset. For each of the models we chose the weights obtained at the number of training steps that gave the best performance on the validation dataset.

## 3. Results

### 3.1. Human detection accuracy compared with CNN performance

The results show that overall for the CNN in both homogeneous & heterogeneous areas we received an F2 score of 0.75. The CNN performed better in heterogeneous areas with an F2 score of 0.778 compared to 0.73 in homogeneous areas. The human annotator median F2 score was 0.78 and performance was better in homogeneous areas – 0.80 compared to 0.766 in heterogeneous areas. These results show that the CNN performed with high comparable accuracy compared to human detection capabilities. Visualisation of one of the model detections is shown in Figure 5.

**Figure 4.**
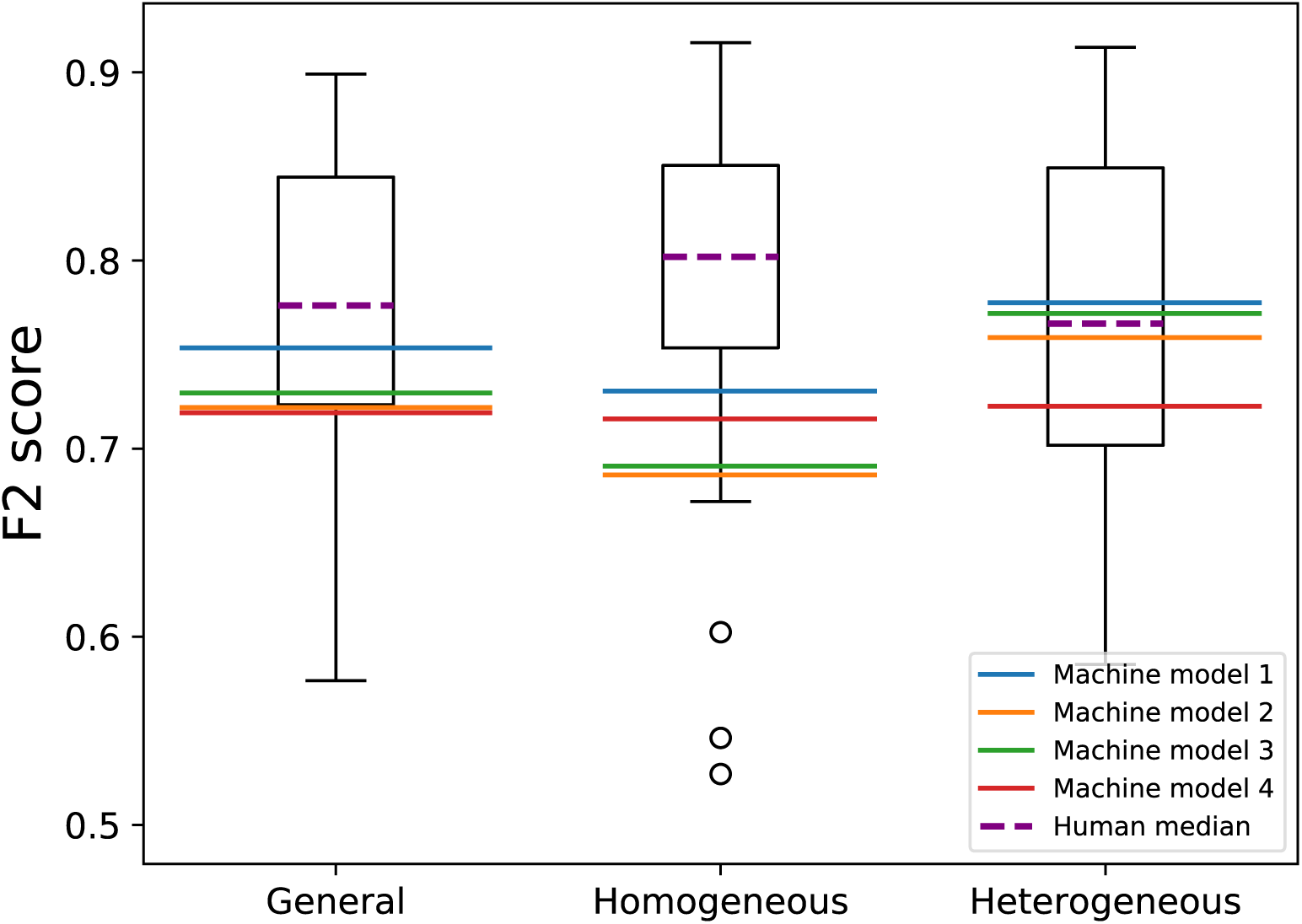
Results of human annotation compared with CNN detection for all images (general) and in homogenous and heterogeneous areas. The boxplots show the human results (circles represent outliers); lines are results from the 4 different CNN models.

**Figure 5.**
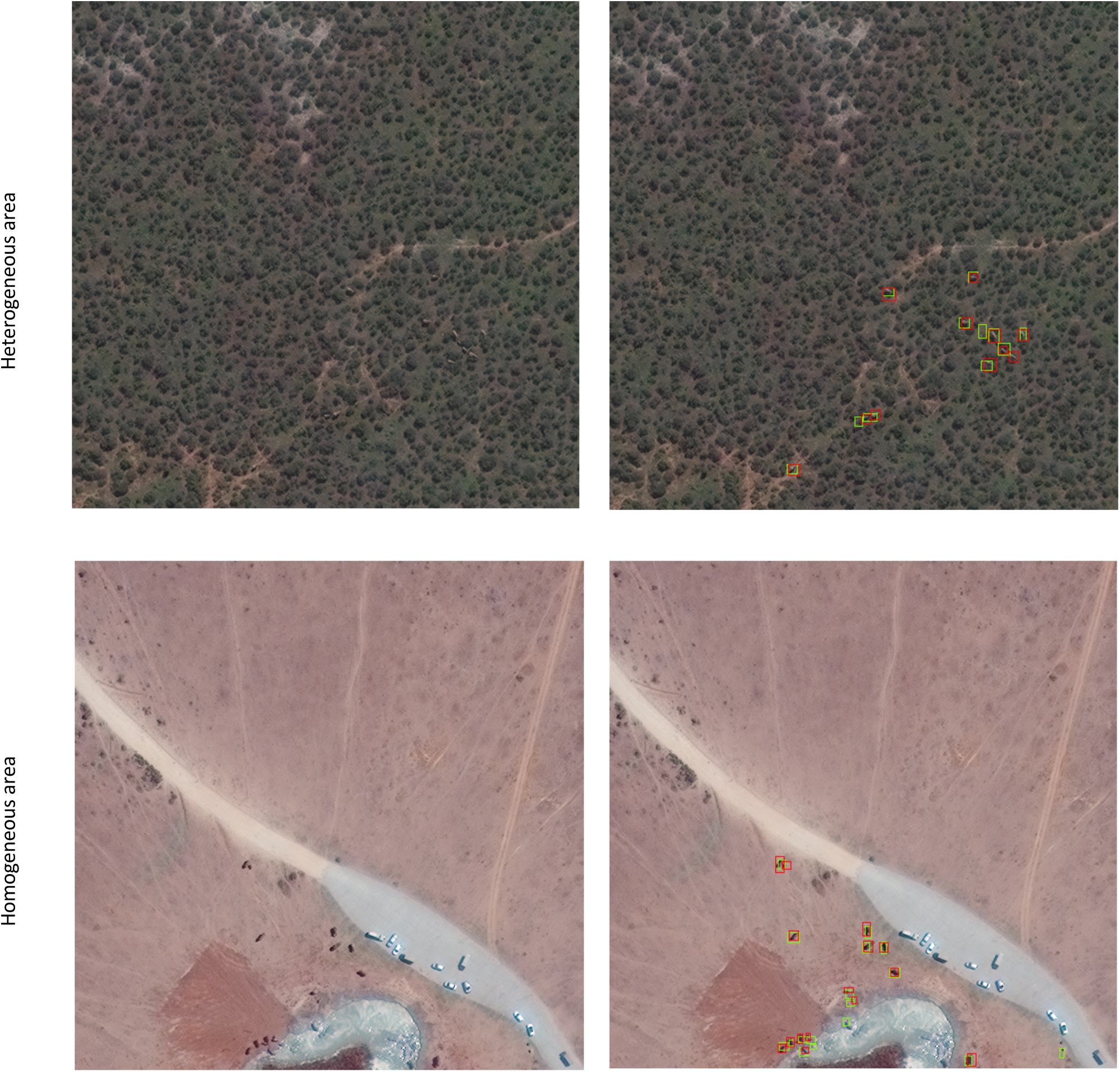
CNN detections: The images on the left are the raw images and images on the right are CNN detections (green boxes) and ground truth labels (red boxes). Satellite image (c) 2020 Maxar Technologies

### 3.2. Testing detection under different image conditions

To test the applicability of the trained CNN model on an elephant population outside of our study area we test, without further training, on a known elephant population in the Maasai Mara in Kenya (Figure 6). The image covers 0.3 km^2^ in which 32 elephants were identified. The CNN managed to detect more than half the elephants in this image (18 true positives) and the resulting F2 score was 0.57. Figure 6 provides visualisation of some example CNN detections.

**Figure 6.**
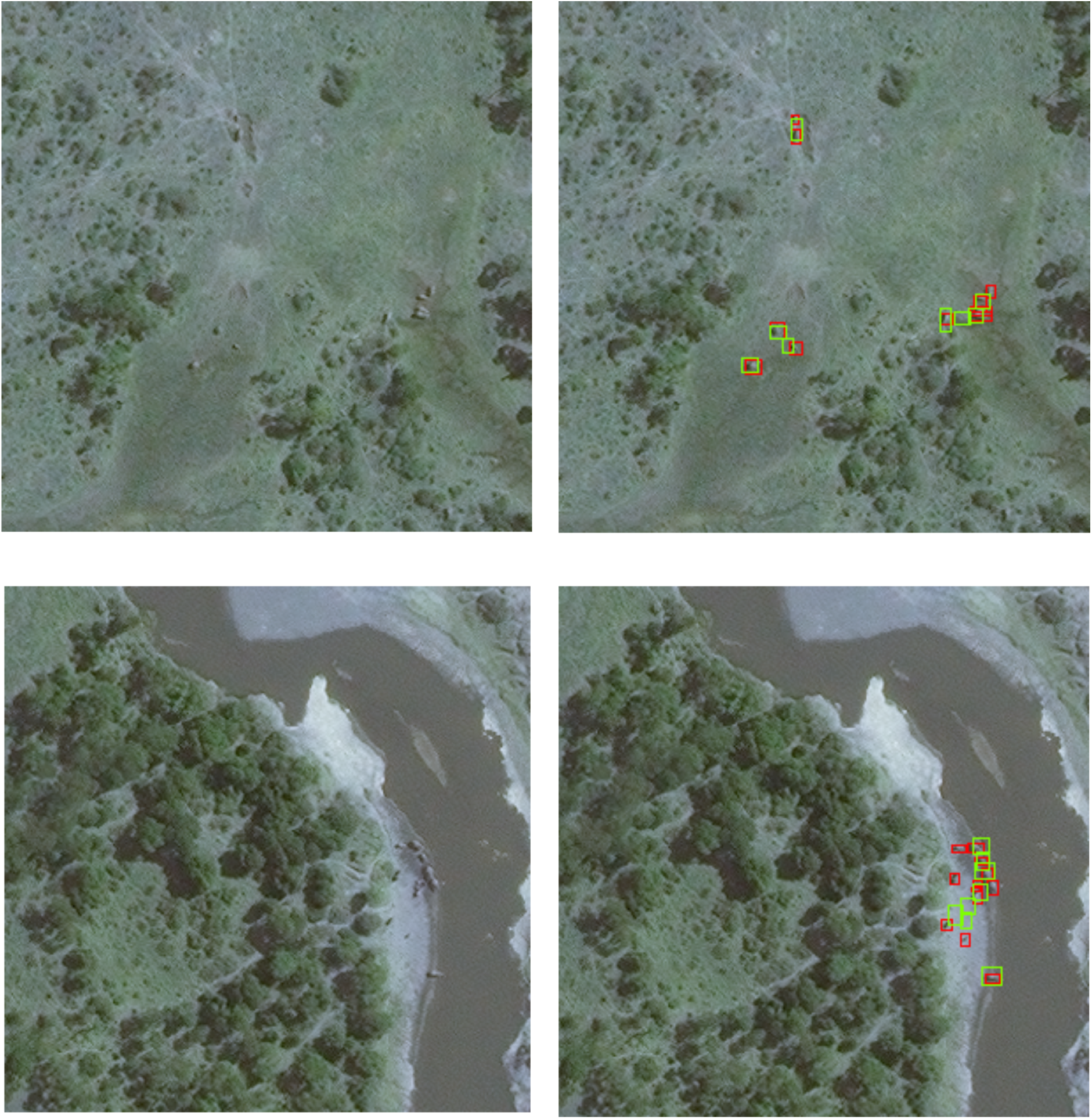
Example of CNN detections in Maasai Mara, Kenya from Geoeye-1 Satellite. Raw images on left and images with CNN detections (green boxes) and ground truth labels (red boxes) on right. Satellite image (c) 2020 Maxar Technologies

## Discussion

For a number of species remote sensing via satellite imagery is already a viable monitoring technique. This study shows the feasibility of this monitoring technique in the case of the African Elephant. From our results we show it is possible to automate detection of African elephants in very high-resolution satellite imagery in both heterogeneous and homogeneous backgrounds using deep learning. We have automated the detection of elephants with as high accuracy as human detection capabilities. There is less variation in the consistency of detection for the CNN compared to human detection performance (as shown in Figure 4). In addition, we show it is possible to generalise detection to elephant populations outside of the site of training data. The transferability of the CNN is promising, as a small amount of training data from this locality or satellite would further increase accuracy. Elephant calves were accurately detected which is noteworthy considering the lack of calves in the training dataset.

Expediating identification of species by automating detection can allow for large-scale application of satellite-based wildlife surveying [26, 46]. The detection process that would usually take weeks can now be completed in a matter of hours. Observer variability mean errors in human-labelled datasets are inconsistently biased while in contrast, false negatives and positives in deep learning algorithms are consistent and can be rectified by systematically improving models. The applicability of this technique is set to increase in the future as the resolution of satellite imagery improves, and costs fall. The new constellation from Maxar, Worldview Legion, will launch in 2021. This constellation has a tropical circle mid-inclined orbit, rather than polar orbit, enabling imagery to be captured in the same location more than 15 times per day at 31cm resolution.

Off-the-shelf object detection tools are increasingly accessible to non-experts; however, the biggest obstacle is obtaining sufficiently large training datasets. Crowdsourced labelling platforms, e.g. Zooniverse [https://www.zooniverse.org/], Amazon Mechanical Turk [https://www.mturk.com/] can help in the creation of these bespoke training datasets using the ‘Wisdom of the crowd’ [81, 82]. A remaining obstacle to scaling this surveying technique is the high cost of commercial satellite imagery. Worldview-3 costs $17.50 per km^2^ for archive imagery and tasking new imagery costs $27.50 per km^2^, with a minimum order of 100 km^2^ (2020 pricing).

Areas of future research to expand this technique include testing whether there are performance improvements for detecting elephants by including the near infrared band and testing for which other species this is already a viable monitoring technique. More broadly, deep learning methods for detecting small objects can be further improved [83, 84] and large training datasets containing images of wildlife from an aerial perspective should be developed. If satellite monitoring is applied at scale then developing methods to ensure standardised and occasional ground-truthing will be required to ensure image interpretation is accurate [25]. Using high resolution satellite imagery as a wildlife surveying tool will inevitably increase in the future as image resolution improves and costs fall. Developing automated detection tools to enable larger scale application of this wildlife monitoring technique is highly valuable as satellite image surveying capabilities expand.

## Acknowledgements

We greatly appreciate the generous support by the DigitalGlobe Foundation (Maxar Technologies) and European Space Agency for awarding the image grants to support this work without which this research would not have been possible. We are also grateful to Hexagon Geospatial for providing a complementary license to ERDAS Imagine which enabled us to process the satellite imagery. We are grateful to all the human volunteer labellers for taking the time to label the images and giving us a point of comparison to the CNN. ID is grateful for a bequest to the Wildlife Conservation Research Unit, University of Oxford from the Ralph Mistler Trust which supported her to carry out this research. SR is grateful for funding from the LLoyd’s Register Foundation through the Alan Turing Institute’s Data Centric Engineering programme. We are grateful to Sofia Minano Gonzalez and Hannah Cubaynes for their valuable comments on the manuscript.

## Data Availability Statement

We are not at liberty to share the raw satellite imagery due to the restrictions defined in the contract with Maxar Technologies and the European Space Agency under which the image grants were awarded.

## Authors’ contributions

I.D. conceived the idea and designed the methodology with oversight from T.W.; I.D. acquired the image data, processed and labelled the satellite images; O.I. conducted all the work in Tensorflow with input from S.R.; I.D. and O.I. organised and analysed the human labels. I.D. led the writing of the manuscript with input from O.I., S.R., D.M. & T.W. All authors contributed critically to the drafts and gave final approval for publication.

